# Arbuscular mycorrhizal fungi, selenium, sulfur, silica-gel and biochar reduce arsenic uptake in plant biomass and improve nutritional quality in *Pisum sativum*

**DOI:** 10.1101/663120

**Authors:** Mohammad Zahangeer Alam, Md. Anamul Hoque, Golam Jalal Ahammed, Lynne Carpenter-Boggs

**Affiliations:** Department of Environmental Science, Faculty of Agriculture, Bangabandhu Sheikh Mujibur Rahman Agricultural University (BSMRAU), Gazipur-1706; Department of Soil Science, Bangladesh Agricultural University (BAU), Mymensingh Bangladesh; College of Forestry, Henan University of Science and Technology, Luoyang 471023, PR China; Department of Crop and Soil Sciences, Washington State University (WSU), Pullman WA 99164-6420 USA

**Keywords:** Arsenic, pea, food chain, AMF, food safety, metal

## Abstract

Arsenic (As) is a carcinogenic substance. It increased in crop grown in field soil from ground water irrigation. Subsequently As transport into the human body through food chains. The reduction of As transport in root, shoot and grain of pea genotypes is significantly important to protect human health. This research is focused on the biomass growth and alleviation of As accumulation in root, shoot and grain of pea genotypes in high As soil (30mgkg^−1^) amended with arbuscular mycorrhizal fungi (AMF), biochar (BC) of rice husk and saw dust, selenium (Se), silica- gel (Si), and sulfur (S). Shoot length, root, shoot and pod mass were generally higher in pea crops grown in soil amended with AMF, Se, Si- gel and S. Rice husk and saw dust BC less consistently increased some growth parameters, particularly in genotype BARI Motor 2. However, the BC’s more often reduced growth and pod mass. All treatments significantly reduced As concentration in tissues; As in grains was reduced on average 60% by any of the soil amendments. AMF, Se and Si- gel all were found more effective than BC for the reduction of As uptake in pea crops. As in grains was reduced 77% by AMF, 71% by Se and 69% by Si- gel on average. As in root, shoot, and grain was also affected by variety; in control treatments, total As uptake in plants pot^−1^ of BARI Motor 1 and 3 was found 60 to 70% higher than BARI Motor 2. Comparing the variety and treatment with most As in grains (BARI Motor 1 control, 0.35 mg As kg^−1^) and least As in grains (BARI motor 1, 2 & 3 with AMF with 0.07 mg As kg^−1^), the choice of variety and soil amendment could reduce human intake of As through pea by 80%. It is recommended that choice of pea variety and soil amendment with AMF and Se have great potential for improving the nutritional quality of pea grown in As contaminated soil, as well as reducing As transfer to human bodies through food chains in pea crops.

## Introduction

Arsenic is a natural toxic element. It has been used as a pesticide, a chemotherapeutic agent and a constituent of consumer products. Arsenic has two forms, inorganic and organic existing in the trivalent or pentavalent state. Trivalent As is generally more toxic than the pentavalent form. A major concern of ingested As is cancer, primarily of skin, bladder, and lung [1]. This metal moves into the human body through the assimilation of food or water. Ground water is the main source for the contamination of As in the soil. It is a global problem, including Bangladesh, Chile, China, Vietnam, Taiwan, India, and the United States [2]. Consequently, As is the number one hazardous substance according to the priority list of the Agency for Toxic Substances and Disease Registry [3].

Simultaneously, Field pea (*Pisum sativum*) is one of the delicious and nutritious pulse crop in Bangladesh as well as throughout the world. It supplies high concentration of antioxidants, starch, protein, fiber, vitamins, minerals and phytochemicals [4]. Fiber in seed coat and cotyledon of pea improves gastrointestinal function. Pea protein, when hydrolyzed, may yield peptides with different bio-active compounds [5]. Among different bio-active stuffs, vitamin and mineral in peas have significant roles for the prevention of different health diseases. Peas also have antioxidant and anti-carcinogenic activities for human beings [6]. However, biomass growth of this crop is being affected due to the contamination of As in soils. Arsenate (AsV) and arsenite (AsIII) both are easily taken up by the cells of the plant root. Once in the cell, AsV can be readily converted to AsIII, which is more toxic. AsV and AsIII both disrupt the plant metabolism of food crops [7]. In pea crops, As inhibited the growth of the roots and shoots (as dry weight) by 65% and 60%, respectively. As well, grain yield (g) and number of pods per plant^−1^ decreased by 66 and 53%, respectively, over controls [8]. Similarly, As decreased biomass growth in wheat and rice crops as well as reduced antioxidant defense activities [9,10,11].

Exposure to As causes substantial stress in crops, including inhibition of growth [12], and physiological disorder [13]. Arsenic acts as a phosphate analogue and is transported across the plasma membrane through phosphate transport systems [12]. Cytoplasmic As(V) interferes on the metabolic processes and cerates toxicity to plants [12,14]. As(III) reacts with sulfhydryl groups (−SH) of enzymes and tissue proteins, inhibiting cellular function, causing death [15]. Even though As is not a redox metal, there is significant evidence that exposure of plants to inorganic As results in the generation of reactive oxygen species (ROS). Reactive oxygen species (ROS) enhance the conversion of As(V) to As(III) in plants [14]. The reactive oxygen species such as, O_2_·^•−^, OH^−^, and hydrogen peroxide enhances oxidative damage [16].

Food adulteration increased in Bangladesh as well as throughout the world due to As toxicity in soils. Pea crops, as well as other food crops, are highly contaminated by As in Bangladesh and West Bengal, India. In this study, several novel methods used for the reduction of As accumulation in root, shoot, and grains of pea genotypes as well as improvement of nutritional quality and biomass growth in As contaminated soils. Biochar (BC), arbuscular mycorrhizas fungi (AMF), selenium (Se), silica –gel (Si- gel) and sulfur (S) were used for the mitigation of As uptake in this crops. BC reduce metalloid uptake in food crops [17,18]. It increases microbial activities as well as reducing As uptake in plants (Gregory et al., 2014). BC is highly potential for reducing As accumulation in spinach crops. The Si-gel and BC both reduces As accumulation in crops as well as releases Si gradually in soils for biomass growth in food crops [19].

Arbuscular mycorrhiza fungi (AMF) can play significant role for increasing plant growth in As contaminated soils. The AMF inoculation reduced As translocation as well as improve biomass growth and nutritional quality of tomato crops [20]. AMF is an effective material for the alleviation of As stress in food crops. AMF may be present in As contaminated soils and are known to exert an ameliorative role on the detrimental effects of As. Although presence of As in soil affects AMF spore germination and colonization. AMF alleviates As toxicity by extending its extra-radical mycelium beyond the depletion zone and helps in nutrient uptake including increase biomass growth of food crops [21]. AMF converts an inorganic As into organic As which is less toxic for plants as well as human beings [22].

However, the presence of As in metal-contaminated soils causing impair of biomass growth in food crops. Selenium (Se) at lower concentration (<1mgkg^−1^) is reported to be stimulatory but is inhibitory at its higher concentration of As in food crops. It is also reported that Se is an effective counter part of As in food crops [23]. An appropriate concentration of Se can be effective against As uptake and translocation in food crops [24]. An antagonistic interaction was reported between Se and As in rice crops. However, subsequent additions of Se helped in mitigating the harmful effects of As and countered the yield reduction caused due to As toxicity. Consequently, the application of Se might be encouraging to reduce As accumulation and toxicity in food crops globally [25].

Similarly, Silicon (Silica-gel) is considered beneficial and valuable element for plant growth, especially under abiotic (metal toxicity) stress [26]. Silicon decrease the concentration of metals uptake in rice such as, Zn, Cd, As and Al [27]. Silicon is considered as an important element for the reduction of metal toxicity in plants. It absorbs heavy metals in roots and minimize their transportation to the shoots [27]. It deposits the SiO_2_ in the apoplast of the leaf surface and roots, which forms a barrier to the flow of metallic ions into grains [28,29]. Arsenic (As) uptake reduce in rice crops through the application of sulfur (S) which is involved in di-sulfide linkage in many proteins and plays a crucial role in As detoxification [30].

Much research has been conducted on the effects of BC, AMF, Se, Si, and S for the reduction of As uptake as well as the improvement of growth, nutrient availability, and bio fortification in different food crops. Effect of soil As on growth and As accumulation in biomass of the BARI released pea genotypes are largely unknown. As is transported from soil to root, shoot, and reallocated to grains in food crops. Subsequently, it is transported into human body through food chains. Therefore, we studied the effect of AMF, Se, BC, Si and Sulfur (S) on growth parameters and As uptake in BARI pea genotypes. It hypothesized that BC, S, Si, Se and AMF will increase plant mass and reduce As concentration as well as improve the nutritional quality of food crops to feed the world.

## Materials and methods

### Background Soil

Background soil was collected from farmer’s field in Bangladesh for growing of pea genotypes in a pot experiment. The soil of the study area is silty loam in the agro-ecological zone of the Old Meghna Estuarine floodplain of Bangladesh, which falls under the order of Inceptisols according to the USDA (United States Department of Agriculture) soil classification. These collected soil samples were brought into the research field of Environmental Science at Bangabandhu Sheikh Mujibur Rahman Agricultural University (BSMRAU). All soil samples were dried under sun light and ground before being used in pots for growing field pea crops.

### Procurement of pea’s seed

Three pea genotypes are developed by Bangladesh Agricultural Research Institute (BARI). Among these genotypes, BARI Motor 1, BARI Motor 2, and BARI Motor 3 pea varieties were procured from Pulse Research Center of BARI. The average yields of these varieties are 10 to 14 tons per ha [31]. Total duration required from seed to maturity is about 80 to 90 days. The production season of these pea genotypes is from November to February in Bangladesh. These varieties were chosen in this study based on their height, life cycle, growing season, and yield.

### Biochars (BC), selenim (Se), silica- gel, sulfer (S), sodium arsenate dibasic heptahydrate (Na_2_HAsO_4_. 7H_2_O), fertilizers, brick pots and pesticides

Raw materials for the preparation of biochars such as sawdust and rice husk were bought from the local market in Bangladesh. Then, the rice husk, and sawdust BC were manufactured using a biochar stove at 400 to 500°C temperature and 3 hrs holding time. This BC is produced by thermal decomposition of biomass under oxygen-limited conditions (pyrolysis). The characteristics of BC are influenced mainly by temperature and biomass. Higher pyrolysis temperature often results in an increased surface area and carbonized fraction of BC, leading to a high sorption capability for pollutants [32]. BC was mixed into soil at 40 g BC kg^−1^ soil. Fertilizers such as Urea, Triple Super Phosphate (TSP), Muirate of Potash (MOP) and Tricho derma-enriched bio-compost were collected as a source of nitrogen (N), phosphorous (P), potassium (K), and other micronutrients. In addition, pots made of brick, fungicides (rubral), and insecticides (chlorpyrifos) were bought from the local market in Bangladesh. On the other hand, selenium metal powder (Se) Qualikems-India, silica gel (Si-gel) Loba chemie- India, sodium arsenate (Na_2_HAsO_4_. 7H_2_O) Sigma-India, and hydrated ferrous sulfate (Fe_2_SO_4_.7H_2_O), Scharlau, Spain were used in this pot experiment.

### Preparation of samples

The collected soil samples from the farmer’s field in Bangladesh were brought into the Department of Environmental Science at Bangabandhu Sheikh Mujibur Rahman Agricultural University (BSMRAU). Initial soil samples of 250-300 (g) were taken from each composite using the guidelines of the [33]. The soil was air dried at room temperature in the laboratory. Samples were then ground and sieved with a ≤250 μm mesh and preserved in polythene bags with proper labeling. Trichoderma enriched bio fertilizers and different BC samples were also prepared for chemical analysis as well as soil samples.

### Analysis of N, P, K, Organic Carbon (OC), and arsenic (As) in background soils

Soil, Trichoderma-enriched bio-fertilizer, and BC’s were analyzed as follows. Total N percentage was determined by the Kjeldhal method [34]. Available P was analyzed by the Olsen method [35]. Exchangeable K was determined by the ammonium acetate extraction method [34]. Organic Carbon (OC) was determined by the wet oxidation method [36]. Sawdust, rice husk BC, Trichoderma-enriched bio-fertilizer, and background soil were digested separately following the heating block digestion procedure for the determination of total As concentration [37] and analyzed by flow injection hydride generation atomic absorption spectrophotometry (FI-HG-AAS, Perkin Elmer A Analyst 400, USA) using external calibration [38] (Table 1).

**Table1.** Total arsenic (As), percentage of OC, N, P, and K in background soil, tricho-derma enriched bio-fertilizer, different biochars (BC) and applied fertilizers

| Samples | Total arsenic (As) in mgkg<sup>-1</sup> | % Organic carbon (OC) | % Nitrogen (N) | % Phosphorus (P) | % Potassium (K) |
| --- | --- | --- | --- | --- | --- |
| Tricho-derma enriched bio-fertilizer | 0.044 | 14.50 | 1.28 | 1.20 | 0.87 |
| Background soil sample | 5.058 | 0.48 | 0.03 | 0.0008 | 0.0016 |
| Saw dust biochar (BC) | 0.003 | 24.60 | 0.33 | 0 | 0.77 |
| Rice husk biochar (BC) | 0.024 | 6.32 | 0.30 | 0.10 | 0.33 |
| Urea | - | - | 46 | - | - |
| Triple Super Phosphate (TSP) | - | - | - | 20 | - |
| Muriate of Potash (MOP) | - | - | - | - | 51 |

### Preparation of high arsenic soil

Collected soil samples were ground uniformly for sowing of the pea seeds in pots. This background soil was measured at a 5.0584 mgkg^−1^ concentration of As. The concentration of As in background soil (5.0584 mgkg^−1^) was increased to 30 mgkg^−1^ through addition of sodium arsenate dibasic heptahydrate (Na_2_HAsO_4_.7H_2_O) as a source of As. Each kg of soil in pots received 0.1041 g sodium arsenate to reach 30 mg As kg^−1^ soil.

### Arbuscular Mycorrhizal Fungus (AMF)

AMF samples were collected from International Culture Collection of (Vesicular) Arbuscular Mycorrhizal Fungi (INVAM), West Virginia University (Morgantown, WV, USA). These AMF samples were found mixed with soils and roots and housed in brick lined pits under the research field of Environmental Science at BSMRAU. A mixture of AMF in soil and roots was cultured in a concrete structured seed bed for multiplication as a source of AMF with the host plant of Sorghum. Before using of AMF in pot soils, Mycorrhizas spores in the soil and vesicle, hyphae, arbuscules in the root samples were observed [39,40]. Pots were inoculated at 40 g AMF soil kg^−1^ pot soil.

### Nutrient augmentation in soils by fertilizers for growing pea crops

Four kg of ground soils with 200g Trichoderma-enriched bio-fertilizers were mixed together in each pot. According to the recommendations of the Bangladesh Agricultural Research Institute (BARI), Urea 90 mg, TSP 180 mg, and MOP 70 mg were incorporated into the soil in each pot. Then, 7–10 pea seeds of each genotype were sowed in each pot.

### Treatments and replications

Three genotypes of BARI released field peas were collected for this pot experiment. These three genotypes of BARI Motor 1, BARI Motor 2, and BARI Motor 3 were selected based on their dissimilar height. With these genotypes there were ten (10) treatments (T_1_ = rice husk biochar, T_2_ = saw dust biochar, T_3_ = AMF, T_4_ = selenium (20 mgkg^−1^), T_5_ = selenium (30 mgkg^−1^), T_6_= silica gel (Si) 5 gkg^−1^ soil, T_7_= silica gel (Si) 10 gkg^−1^ soil, T_8_= sulfur (S) Fe_2_So_4_.7H_2_O 50 mgkg^−1^(S), T_9_= sulfur (S) Fe_2_SO_4_.7H_2_O 100 mgkg^−1^ (S), and T_10_ = control). All soils were prepared at an arsenic concentration of 30 mgkg^−1^. Five replications were used in this pot experiment and total number of pots was 150. As a final point, these three field pea genotypes were also grown in 5.0584 mgkg^−1^ As concentrate background soils in pots. Five replications also followed in this stage for a total number of 15 pots.

### Shoot length, dry weight of root, shoot, and pods, grinding and sieving of samples

At random, average shoot lengths of pea crops were measured using a measuring tape (cm) at week 10 in each treated pots. The average dry weight of the root, shoot and pods of the pea crops was measured separately using electrical balance (g) after harvesting in each As treated pot during week 14. All samples were dried in an oven at 55°C for 72 hours towards the digestion of samples for the determination of As uptake in root, shoot and grains of pea crops. Grains were separated from pod by hand with gloves. Gloves in hand were changed during the separation of grain for each samples. Then samples were ground separately by coffee grinder using liquid nitrogen. This grinder was cleaned between the samples through tissue paper with ethyl alcohol (C2H5OH). These ground root, shoot and grain samples were sieved with 250μ mesh. Then all samples were kept in envelopes with proper labeling.

### Digestion of samples

Soils, roots, shoots and grains of pea crops were digested separately following the heating block digestion procedure [37]. Of the soil samples including tricho-derma and bio char’s, 0.2 g were taken into clean, dry digestion tubes and 5 ml of concentrate HNO_3_ and 3 ml concentrate HClO_4_ added to it. The mixture was allowed to stand overnight under a fume hood. On the following day, this vessel was put into a digestion block for 4 hours at 120° C temperature. Similarly, 0.2 g ground root, shoot and grain samples were put into clean a digestion vessel and 5 ml concentrate HNO_3_ was added to it. The mixture was allowed to stand overnight under the fume hood. On the following day, this vessel was put into the digestion block for 1 hour at 120° C temperature. The content cooled and 3 ml HClO_4_ was added to it. Again, samples were put into the heating block for 3-4 hours at 140°C. Generally heating stopped whenever a white dense fume of HClO_4_ was emitted into air. Then samples were cooled, diluted to 25ml with de-ionized water and filtered through Whatman No 42 filter paper for soil and plant samples. Finally, samples were stored with polyethylene bottles. Prior to sample digestion, all glassware was washed with 2% HNO_3_ followed by rinsing with de-ionized water and drying.

### Analysis of total arsenic

Digested samples were brought into the laboratory of chemistry at Bangladesh Atomic Energy Center, Dhaka for the analysis of total As in the pea root, shoot, and grain samples. The total As in root, shoot, and grain of pea crops were analyzed by flow injection hydride generation atomic absorption spectrophotometry (FI-HG-AAS, Perkin Elmer A Analyst 400, USA) using external calibration [38]. The optimum HCl concentration was 10% v/v and 0.4% NaBH_4_ produced the maximum sensitivity. Three replicates were taken from each digested sample and the mean values obtained based on the calculation of those three replicates. Standard Reference Materials (SRM) from National Institute of Standards and Technology (NIST), USA analyzed by the same procedure at the start, during and at the end of the measurements to ensure continued accuracy.

### Statistical Analysis

The design of this experiment followed the Completely Randomized Design (CRD). Analysis of Variance (ANOVA), and mean comparison of treatment effects on the reduction of arsenic uptake in root, shoot and grain of pea crops were analyzed using Software R.

## Results

### Shoot length

The length of shoot of BARI Motor 1 & 2 pea genotypes grown in soils with an arsenic concentration of 30 mgkg^−1^ treated with arbuscular mycorrhizal fungi (AMF), biochars (BC), selenium (Se), sulfur (S) and silica (Si-gel) were found statistically similar. The shoot length of BARI motor 1 treated with AMF, Se and S was found significantly higher (*p≤ 0.001*) than control. However, the shoot length of BARI motor 2 treated with Se (T_5_) and S (T_8_) both were found significantly higher than control during the week 10 at 30 mg As kg^−1^ soils. Selenium (Se), Si-gel and S treated shoot length of BARI Motor 3 pea genotypes grown in soils with an As concentration of 30 mgkg^−1^ were found significantly higher than control at week 10. Treatment of AMF was found statistically similar with the treatment of Si-gel and S for increasing shoot length in BARI Motor 3 pea genotypes during the week 10 at 30 mgAskg^−1^ soils. The shoot length of these pea genotypes increased by 8, 20, 31, 28, and 20% following BC, AMF, Se, S, and Si- gel treatments, respectively. Notably, Se is possibly effective for increasing of the length of shoot in pea genotypes grown in soils with an As concentration of 30 mgkg^−1^ (Fig 1).

**Fig 1.**
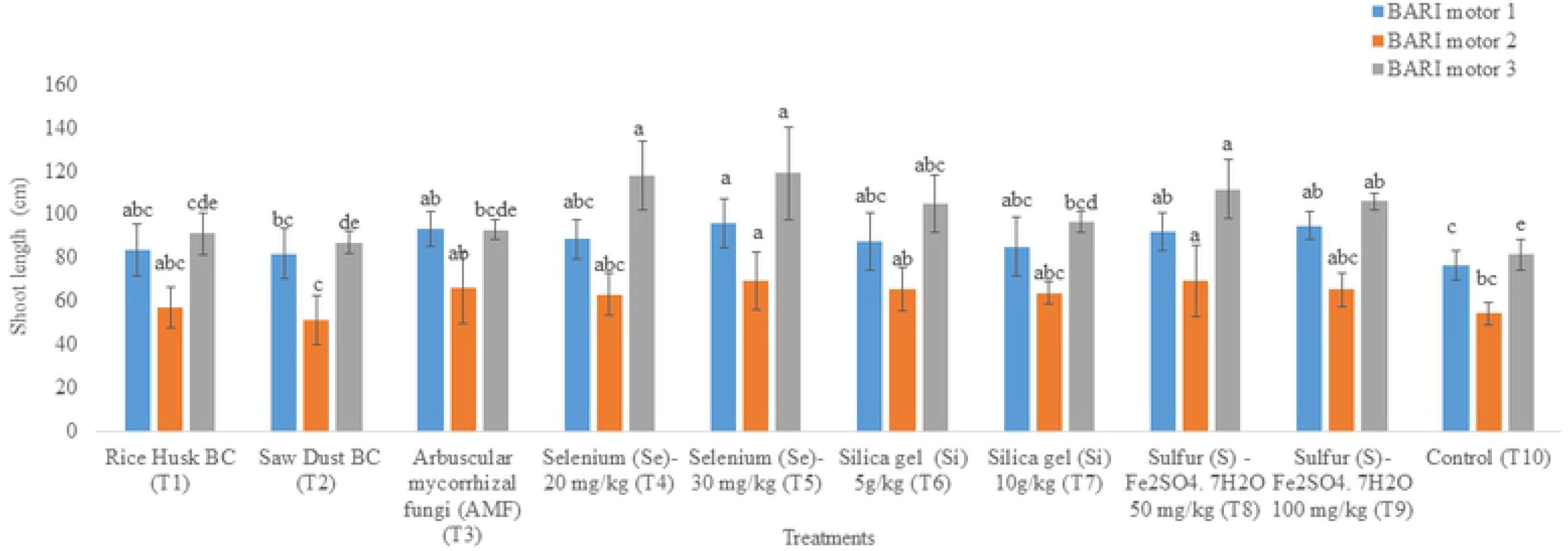
Effect of Biochar (BC), AMF, Se, Si-gel and sulfur (S) on shoot length (Mean ± SEM) of BARI Motor 1, 2 & 3. pea genotypes grown in soils with an arsenic concentration of 30 mgkg^−1^ at week 10. Means denoted by different letters under the same arsenic level indicate significant difference at 0.1% level of significance.

### Root, shoot and pod mass of pea genotypes

In BARI Motor 1, 2 &3 pea varieties in 30 mgAskg^−1^ soil, root biomass was found statistically similar among the treatment of AMF, BC, Se, Si-gel, and S. However, Se (T_5_) and BC (T_1_) significantly increased the root mass in BARI Motor 1 pea genotypes compared with that of control. As well, the root mass of S treated (T_9_) BARI Motor 3 was found statistically higher (*p*≤*0.001*) than that of control at week 14. The root mass of these pea genotypes increased by 30%, 52%, 55%, 52%, and 33% following BC, AMF, Se, S, and Si- gel treatments, respectively. AMF, Se and S are equally effective for increasing root mass in these pea genotypes (Fig 2). The shoot mass of AMF, Se, Si-gel, and S treated BARI Motor 1 & 3 were found significantly higher than that of control during the week 14 at 30 mgAskg^−1^soils. In BARI Motor 2, all treated shoot mass was found statistically similar with control at same concentration of As in soils. The shoot mass of pea genotypes increased by 13%, 70%, 92%, 92% and 80% following BC, AMF, Se. Si-gel and S treatments, respectively (Fig 3). The pod mass of AMF, Se, Si gel and S treated BARI Motor 1 &3 were significantly increased compared with that of control during the week 14 at 30 mgAskg^−1^soils. AMF treated pod mass in BARI Motor 2 was found statistically similar with Se, Si-gel, BC, and S treated pod mass. The pod mass of these pea genotypes increased by 45%, 54%, 45% and 45% following AMF, Se, Si gel, and S treatments, respectively (Fig 4). Dry weight of root, shoot and pod were found to be significantly higher in BARI Motor 1 & 3 than BARI Motor 2 pea genotypes.

**Fig 2.**
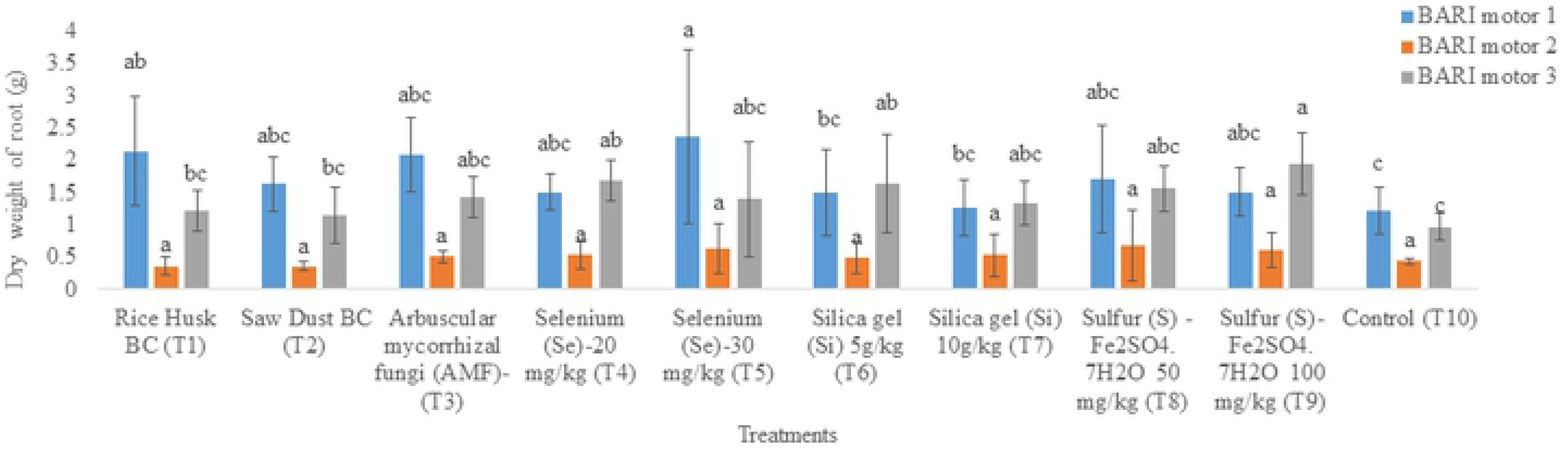
Effect of Biochar (BC), AMF, Se, Si-gel and sulfur (S) on root mass (Mean ± SEM) of BARI Motor 1, 2 & 3. pea genotypes grown in soils with an arsenic concentration of 30 mgkg^−1^ at week 14. Means denoted by different letters under the same arsenic level indicate significant difference at 0.1% level of significance.

**Fig 3.**
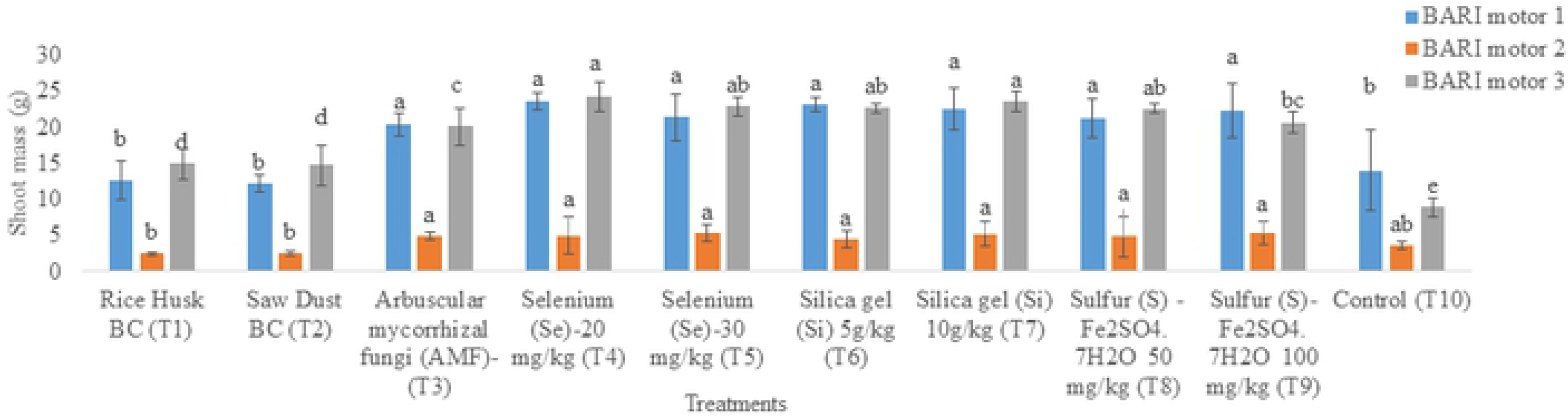
Effect of Biochar (BC), AMF, Se, Si-gel and sulfur (S) on shoot mass (Mean ± SEM) of BARI Motor 1, 2 & 3. pea genotypes grown in soils with an arsenic concentration of 30 mgkg^−1^ at week 14. Means denoted by different letters under the same arsenic level indicate significant difference at 0.1% level of significance.

**Fig 4.**
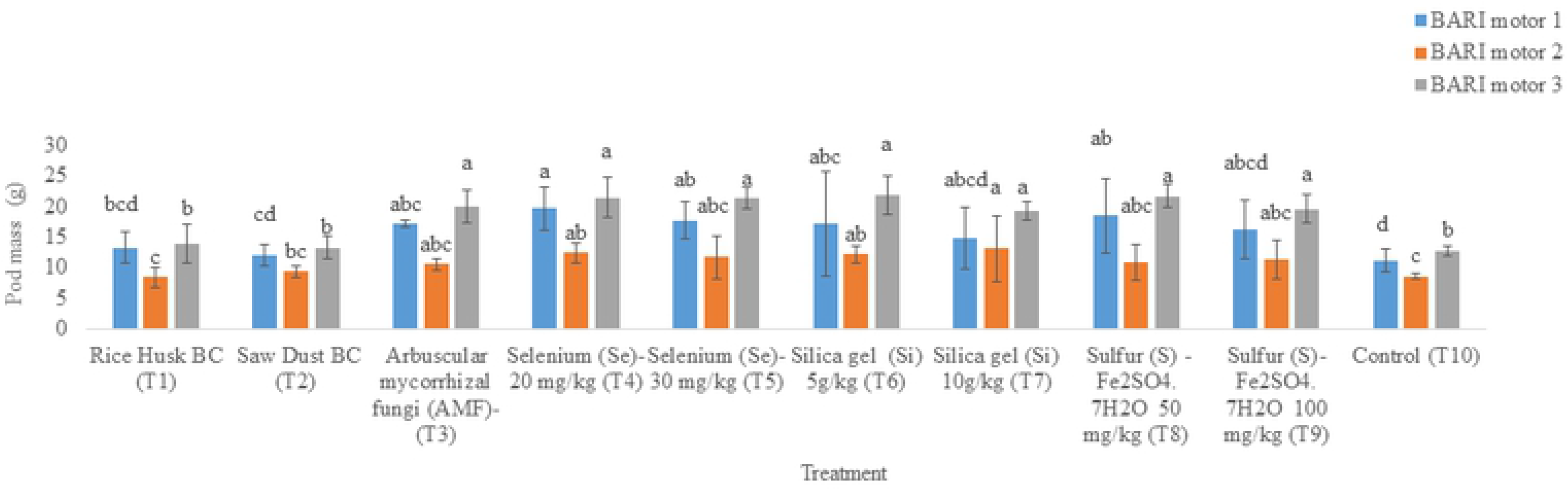
Effect of Biochar (BC), AMF, Se, Si-gel and sulfur (S) on pod mass (Mean ± SEM) of BARI Motor 1, 2 & 3. pea genotypes grown in soils with an arsenic concentration of 30 mgkg^−1^ at week 14. Means denoted by different letters under the same arsenic level indicate significant difference at 0.1% level of significance.

### Treatments with BC, AMF, Se, Si-gel and S reduced arsenic uptake in root, shoot and grain in pea genotypes

According to ANOVA, treatment and interaction of variety & treatment significantly affected As concentration in root, shoot and grain in BARI Motor 1, 2 & 3 pea genotypes grown in soils with an As concentration of 30 mgkg^−1^ (*p*≤*0.001*;*p*≤*0.0001*;*p*≤*0.0005*) (Table 2). Treatments with BC, AMF, Se, Si-gel and S significantly reduced As concentration in the root of pea genotypes as compared to control. Similarly, all treatments had significant effect on the reduction of As concentration in shoot and grain of pea genotypes as compared to control. AMF, Se and Si-gel were found highly effective for the reduction of As concentration in root, shoot and grain of pea genotypes grown in soils with an As concentration of 30 mgkg^−1^ in comparison with BC, S and control. As in grains was reduced on average 53% by BC soil amendments. Likewise, As in grains was reduced (on an average) 77%, 71%, 69% and 66% by AMF, Se, Si-gel, and S treatments, respectively. AMF, Se, Si- gel and S all were found more effective than BC for the alleviation of As concentration to grain in pea crops (Table 3). Similar concentration of As was found in grain of BARI Motor 2 and 3 pea genotypes grown in background soils (As concentration 5.0584 mgkg^−1^) (*p*≤ *0.05*) (Table 3).

**Table 2.**
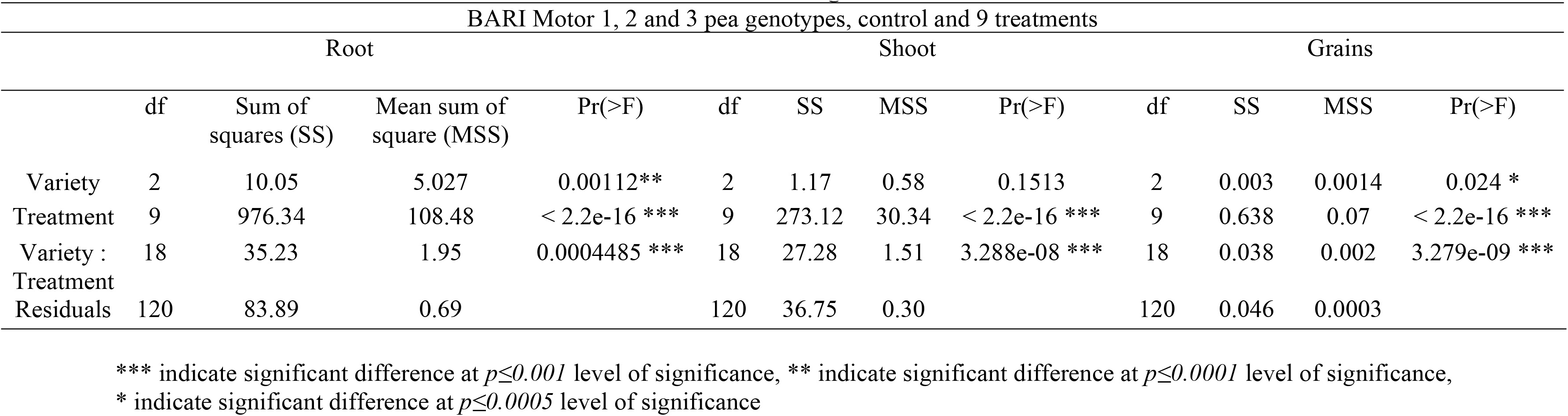
ANOVA on the concentration of arsenic in root, shoot and grains

**Table 3.**
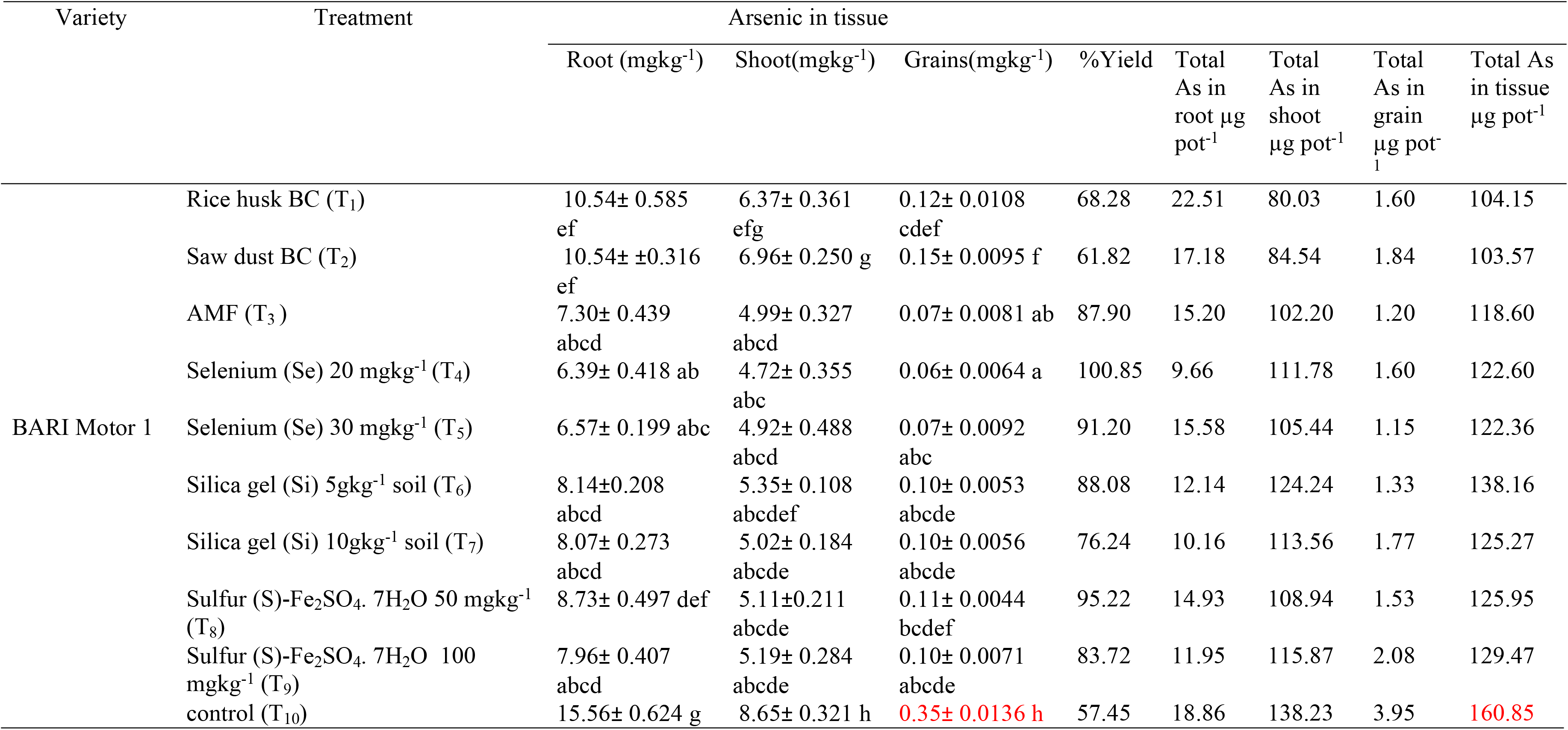
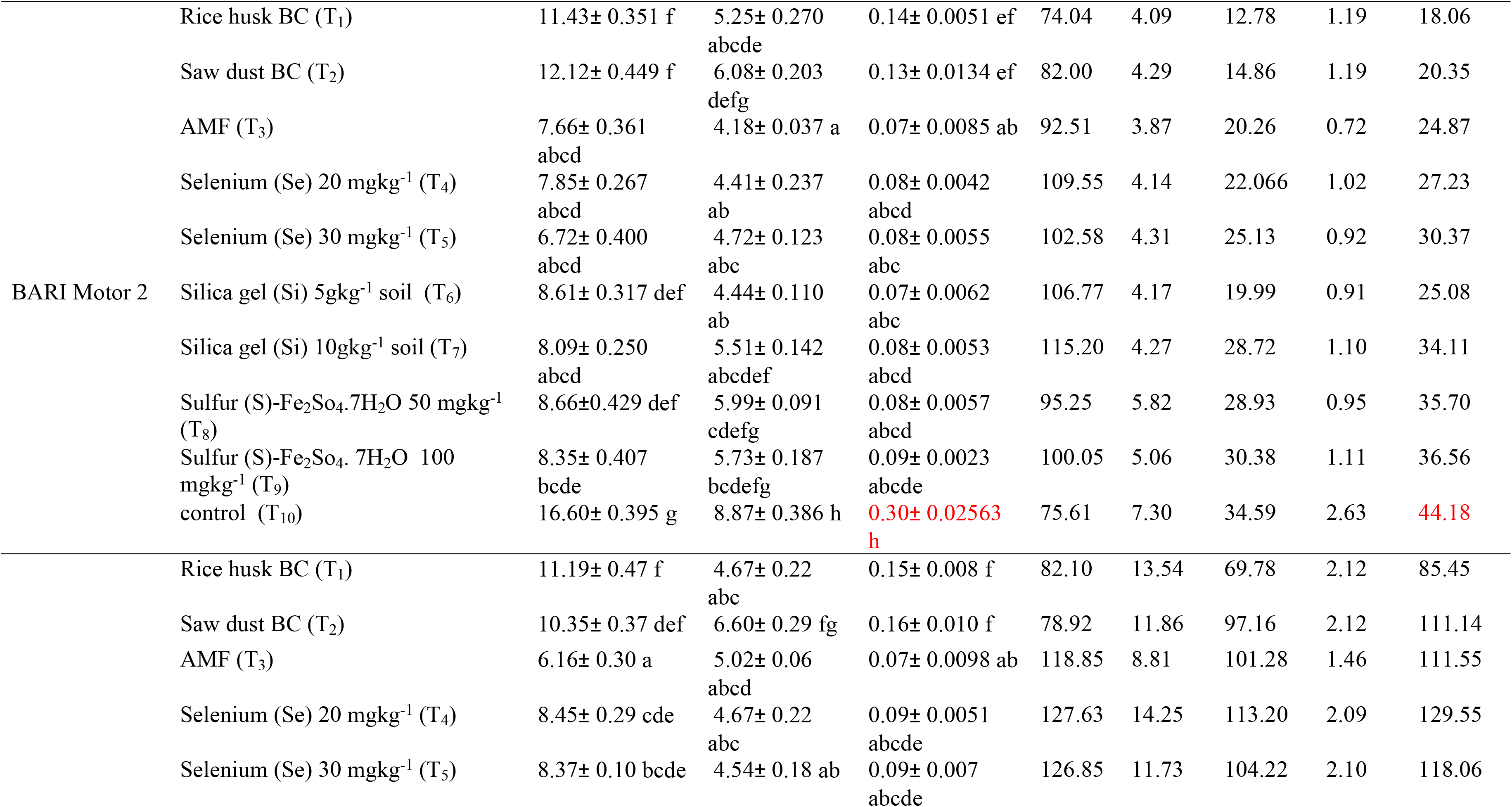
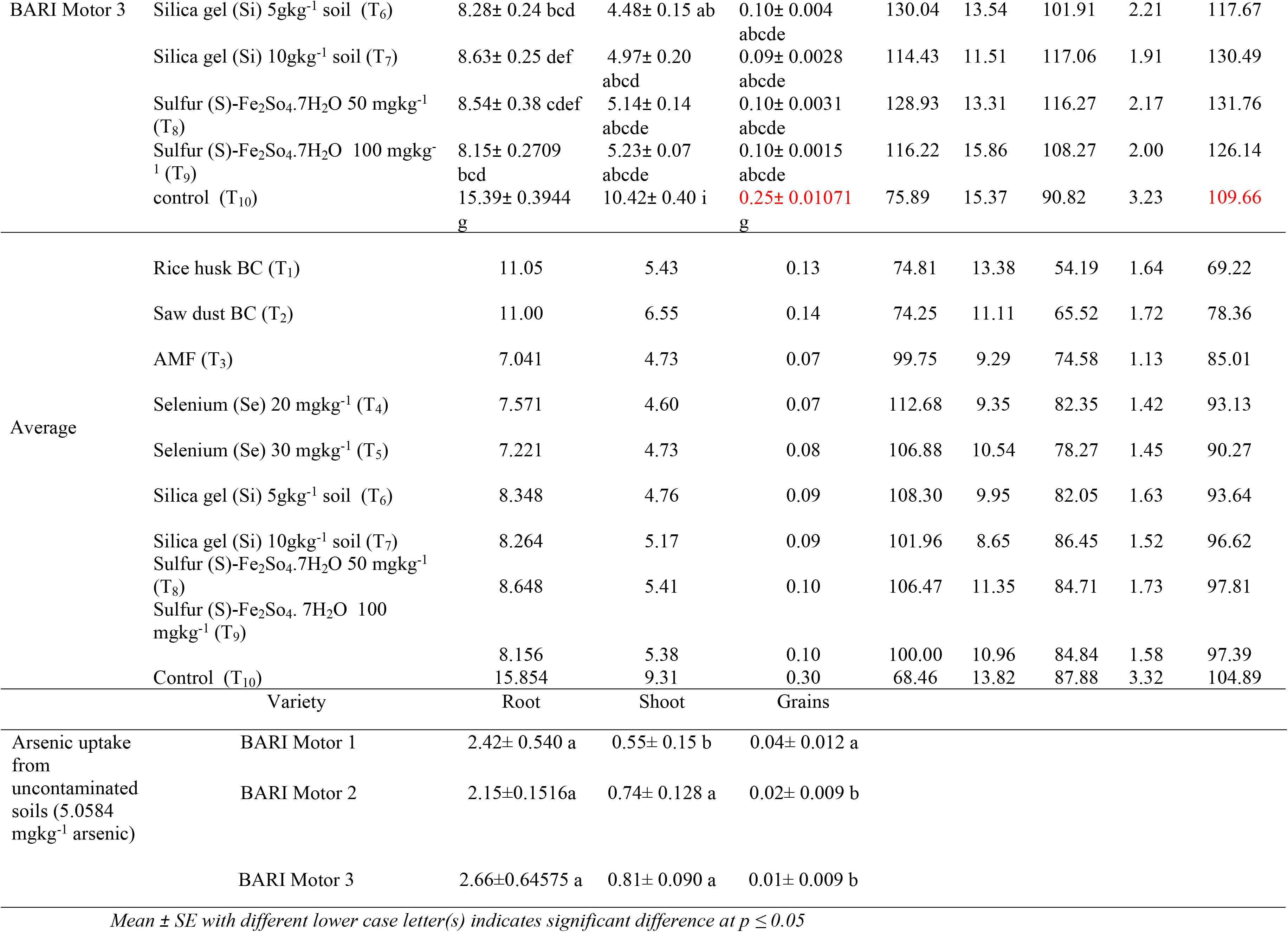
Arsenic concentration in root, shoot and grains of BARI Motor 1, 2 & 3 pea genotypes grown in soils with an arsenic concentration of 30 mgkg^−1^

## Discussion

Arsenic (As) is lethal to all life forms and is a leading carcinogen. Its accumulation in food crops and subsequent ingestion pose a severe risk to public health worldwide [41]. Due to the extensive use of groundwater for irrigation in Bangladesh and west Bengal of India, As toxicity has increased in surface soil from its sources. Arsenic deters the biomass growth such as, root, shoot and pod mass of food crops including pea genotypes. It creates a health hazards to human beings through food chains as well [42]. For this reason, biochar (BC), arbuscular mycorrhizal fungi (AMF), selenium (Se), silica gel (Si-gel), and sulfur (S) were applied for the improvement of biomass growth as well as reduction of As concentration in root, shoot and grain of pea crops grown in soils with an As contamination of 30 mgkg^−1^.

BC enhance microbial activities as well as increase biomass growth under metalloid stress conditions [43]. Oxidative stress induced due to the contamination of As in soils. BC reduce oxidative stress under As stress condition as well as increase the biomass growth of wheat crops [44]. In garden pea, BC increases root and shoot mass and grain yield as well as enhancing the percentages of germination in abiotic stress condition [45]. Similarly, shoot length including root, shoot and pod mass of pea genotypes grown in soils with an As concentration of 30 mgkg^−1^ had increased significantly in BC applied soils under As stress conditions as compared to control (Figs.1–4). BC increase cation exchange capacity (CEC), and availability of soil macro- and microelements in As contaminated soils [46,47,48,49]. In addition, BC reduce mobility of other heavy metals through altering redox potential in tomato and sweet corn crops [18,50,51]. Likewise, BC reduce the transportation of As in root, shoot and grain of pea genotypes grown in soils with an As contamination of 30mgkg^−1^ as compared to the control (Table 3).

Aurbuscular mycorrhizal fungi (AMF) increases root and shoot length along with enhances dry matter of root shoot and pod in pea crops at 30 mgkg^−1^ As concentrated soils. Consequently, biomass production was found higher with AMF treated pea genotypes than control (Figs 1–4). Similarly, AMF effectively reduced As concentration in grain of lentil crops as well as improve the biomass growth and nutritional quality [52]. Much research was conducted on the effect of AMF on biomass production and antioxidant activities in wheat crops under abiotic stress [53]. AMF generally consists of hyphae, arbuscules, spore, mycelia and vesicle with the mixture of soil around the root of host plant. Due to the hyphal network with host plant in the rhizosphere; they can promote plant growth, yield and quality of crops [54,55]. In this circumstances, AMF-induced positive effects on biomass growth as well as improving nutritional quality in different plant species including, pea, kidney bean, pepper, watermelon, muskmelon, onion, tomato and asparagus [55, 56,57,58,59].

AMF and rhizobia both are compatible between each other with the host plant, can eventually increase the growth, yield and nutritional quality of legume crops [60]. In addition, AMF inoculation could effectively improve growth of cucumber including different vegetable crops [58]. AMF species such as *Funneliformis mosseae* BEG167, *Rhizophagus intraradices* BEG141, and *Glomus versiforme* Berch enhances photosynthetic pigments, and antioxidant activities in food crops [52, 61, 62] reports AMF of *Glomus versiforme* and *Rhizophagus irregularis* both enhances biomass growth, gas exchange and chlorophyll fluorescence in black locust seedlings.

The rate of As accumulation in root, shoot and grain of AMF treated pea crops grown in soils with an As concentration of 30 mgkg^−1^ was found similar to the uncontaminated soils (Table 3). Mycorrhizal plants display lower specific As(V) uptake and higher P: As ratio than non-mycorrhizal plants [63]. As (III) enters in the plants through phosphate transporters as a phosphate analog or through aquaglyceroporins [64]. Detoxification mechanisms for As (III) include efflux from the roots, sequestration in cell vacuoles and complexation with thiols for which As(III) has very high-affinity [65]. Food crops that are adapted to As-polluted soils are generally associated with AM fungi [66, 67]. Inoculation by AM fungi can exert protective effects on vascular plants under As contamination by transforming inorganic As in less toxic organic forms or by diluting As concentration by enhancing plant biomass [22, 59, 67, 68, 69, 70]. Likewise, As accumulation in root, shoot and grain of pea crops had remarkably reduced through the application AMF in this study (Table 3).

Selenium (Se), Si- gel and sulfur (S) all are significantly increases the biomass growth of pea crops grown in soil with an As concentration of 30 mgkg^−1^ (Figs 1–4 and Table 3). Se is an essential element for plants benefits as well as human beings. Much research has been validated on the benefits of Se with respect to the productivity of certain vascular plants [71,72,73,74]. Se translocated into grains as well as increases yield and antioxidant activity in food crops [71,73,75]. Likewise, Se application increases biomass growth and yield of pumpkin (*Cucurbita pepo* L.), lettuce (*Lactuca sativa* L.) and canola (*Brassica napus* L.) crops as well as enhances photosynthetic pigments and antioxidant activities under abiotic stress [52,74,76,77,78]. Similarly, Se increase biomass growth in BARI released pea genotypes in this study (Figs 1–4). In addition, it moves into human bodies through grain which is significantly important for the reduction of Se deficiency in humans [79]. As well, Se application in the As contaminated environment reduce As uptake and phyto-toxicity by modulation of phenolic compounds and increase biomass growth and nutritional quality of food crops [80,81].

Se and Si both reduce metal uptake in biomass as well as enhances antioxidant activities and biomass growth under heavy metal stress in food crops [82]. In rice crops, Si enhances yield components such as, number of spikelet’s per panicle and the percentage of filled spikelet’s, [83,84]. Likewise, silicon (Si) addition reduces the accumulation of Cd and As in the edible parts of potato, carrot, onion, and wheat plants [85]. Also Si increased biomass growth in sunflower and lettuce crops in As stress condition as well as reduce As uptake in the edible part of these crops [86,87]. Transporter of Si formed nodulin-26 like intrinsic proteins (NIPs) with plants grown in As contaminated soils [88]. Si competes with arsenite [89] and reduce arsenite uptake in biomass of food crops from contaminated soils [90]. This interaction (Si & As) has received much attention in recent years [91,92,93] for mitigating As toxicity in food crops. Similarly, As uptake significantly reduces Si treated pea crops grown in soils with an As concentration of 30 mgkg-^1^(Table 3).

Application of Sulfur (S) in As contaminated soils increase biomass growth in plants [94]. S reduce As translocation from soils to roots and grains in legume crops. This element (S) induced iron plaque and glutathione in leaves and roots that mechanism deters As uptake in the biomass of food crops [95]. Arsenic uptake hindered by the application of S content fertilizers in rice plants, however, variable results have been observed [96,97,98]. A recent study by [99] assessed the effect of S (sulfur) on As accumulation and distribution in rice (*Oryza sativa*) plants and found As accumulation at zero in comparison to control. Availability of S (sulfur) in soils with an As stress in food crops rely on thiol metabolism. As a result, high doses of S reduce As concentration in grain (44%) in comparison to control [97].

Total As uptake by plants was reduced by most soil treatments in most varieties (Table 3). This is true despite the great increase in tissue and pod yields due to Se and AMF treatments. BC’s on average reduced total As uptake by 30%, and Se and AMF by 13% and 19%, respectively (Table 3). The larger reduction in total uptake by BC’s is largely due to the loss of biomass in BC treatments, vs. increase in biomass in Se and AMF treatments. Concentration of As in roots was consistently reduced by AMF and Se, giving no indication that As was sequestered within the root or AMF tissue. Proportion of total plant As uptake held in roots was also found lower in AMF and Se treated crops. If AMF were reducing translocation of As from roots into shoots, concentration or proportion of As in roots of mycorrhizal plants would be greater than that in non-mycorrhizal plants. Rather, the proportion of total As uptake found in shoots vs. pods was affected by both Se and AMF but not by BC’s. The proportion of As uptake found in shoots was increased 6% by Se and 4% by AMF, and decreased in pods by 49% by Se and 50% by AMF. Choice of pea variety should be determined by the soil condition. On average, these 3 varieties yielded only 97% as much pod mass in high As soil as in low As soil (Table 3). AMF and Se treatments improved pod yields in high As soils of these pea varieties.

Arsenic induced oxidative stress in food crops. As toxicity includes the As-induced ROS reactions with macromolecules: lipid peroxidation and protein and nucleic acid damage. As a result, reduce enzymatic antioxidant defense system and mobilize the cell to synthesize low-molecular-weight antioxidants which are important in the prevention of ROS-induced damage [100]. As-induced reactive oxygen species (ROS) impacts on plants at biochemical, genetic, and molecular levels. Different enzymatic (superoxide dismutase, catalase, glutathione reductase, and ascorbate peroxidase) and non-enzymatic (salicylic acid, proline, phytochelatins, glutathione, nitric oxide, and phosphorous) activities declined through the contamination of As soils in plants [101]. However, future agriculture will depend on food security as well as providing of contaminant free food throughout the world. For this reason, phytoremediation of As accumulation in edible parts of pea crops using AMF, BC, Se and Si –S are significantly important for the upcoming demand in agriculture all over the world. Furthermore, these As relief substances (AMF, Se, Si, BC and S) reduced oxidative damage and osmotic stress as well as increased biomass growth and nutritional quality in food crops [11,102].

## Conclusion

Arsenic is the number one hazardous substance in the world. It is transported from soils to root, shoot, and grains in different food crops. Among various food crops, pea provides essential vitamins and minerals to human beings. However, biomass growth and yield reduces in crops grown in As contaminated soils. Subsequently this metalloid is transported into human bodies through the consumption of contaminated foods. In this situation, bio-char (BC), arbuscular mycorrhizal fungi (AMF), selenium (Se), silica- gel (Si), and sulfur (S) were used for the reduction of As concentration in root, shoot, and grain of pea genotypes. AMF, Se and Si- gel all were found more effective than BC for the alleviation of As concentration in tissues. As in grains was reduced 77% by AMF, 71% by Se and 69% by Si- gel on average. These As relief substances reduces oxidative damages as well as increases biomass growth and nutritional quality of pea genotypes grown in soils with an As concentration of 30 mgkg^−1^. If similar results are found in subsequent field studies, promotion of BARI pea crops with AMF could increase yields in high As fields by over 100%, while decreasing concentration of As in the pea by 76%. Because AMF colonization in field soils greatly increase the effective rooting volume, water availability, and nutrient uptake, the effect of AMF in increasing yield and reducing As uptake in field soils may be even greater in field soils than in a pot study. This relatively simple change would provide a meaningful improvement in food quantity, quality, and security for communities’ dependent on high As soils.

## Acknowledgement

The authors are grateful to the laboratory of Crop and Soil Sciences at Washington State University (WSU), WA, USA and the Laboratory of Environmental Science at BSMRAU. This research is also recognized as part of a collaborative PhD program between Washington State University and the Bangladesh Agricultural University (BAU): Mymensingh-2202. Authors are especially grateful to Rebecca McGee (WSU-USDA scientist), Tarah S. Sullivan (WSU) and Shah Mohammad Naimul Islam (A/Prof, BSMRAU) for their conceptualization and statistical analysis on the development of research plan.

## Supporting Information

**Dataset. AMF, BC, Se, Si-gel and S reduced total As concentration in plant biomass pot^−1^**

